## Supplemental Figure S1 for "Intracellular conformation of amyotrophic lateral sclerosis-causative TDP-43"

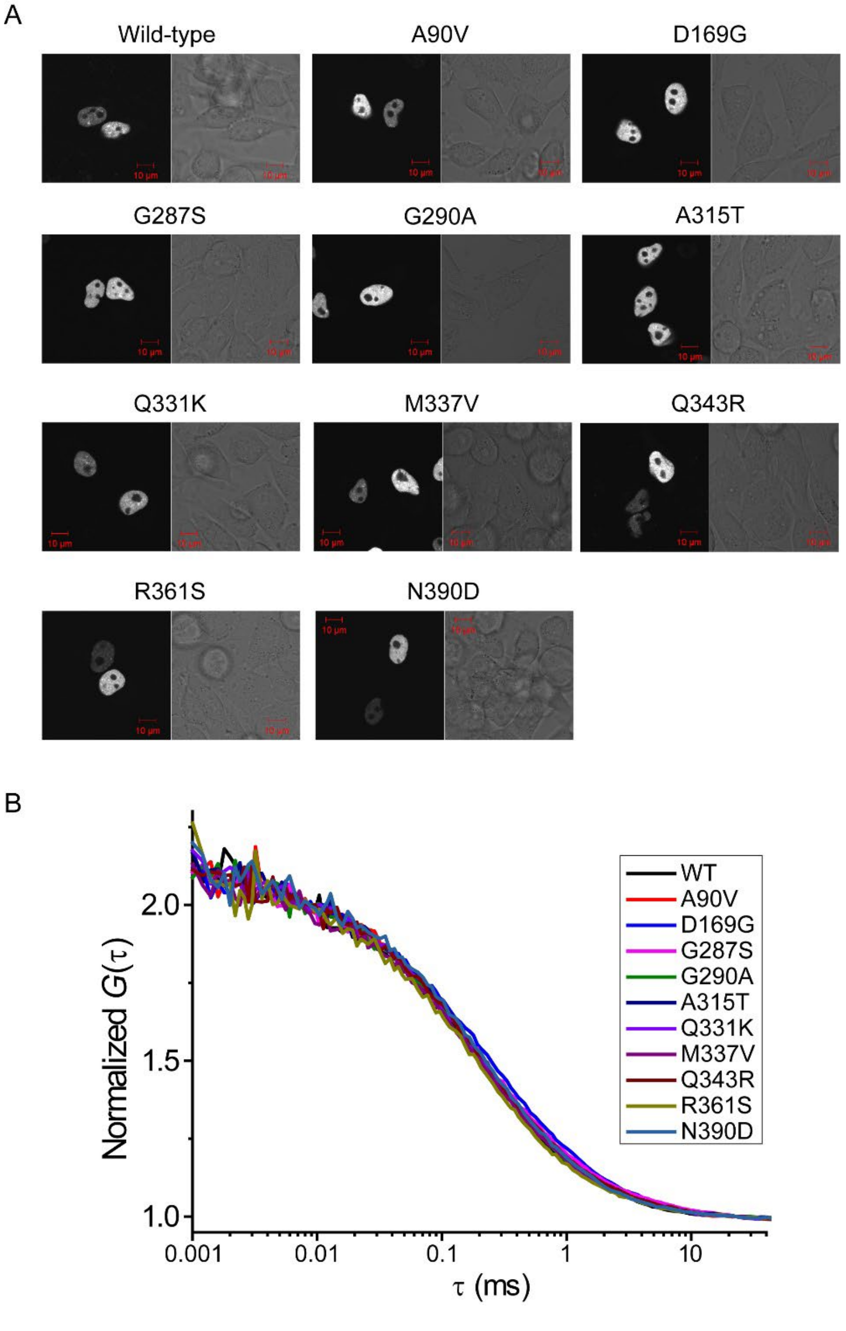


**Supplemental Figure S1 Subcellular localization and diffusion state of ALS-associated mutants of TDP-43 tagged with eGFP.**

(A) Confocal fluorescence images of eGFP-labeled wild-type and ALS-associated mutants of TDP-43. Alphabets and numbers indicate the one-letter symbol of the amino acid containing the mutation and the location of the amino acid residue. Bars = 10 µm. (B) Normalized autocorrelation functions of eGFP-labeled wild-type (WT) and ALS-associated mutants of TDP-43 in cell lysates. No significant differences in the autocorrelation curve were observed.
